# Rewiring DNA Repair Activity into CRISPR Signal Transduction via Synthetic DNA Transducers

**DOI:** 10.1101/2025.07.11.664310

**Authors:** Neda Bagheri, Alessandro Bertucci, Rosa Merlo, Alessandro Porchetta

## Abstract

Harnessing the precision of CRISPR systems for diagnostics has transformed nucleic acid detection. However, the integration of upstream cellular signals into CRISPR-based circuits remains largely underexplored. Here, we introduce a synthetic transduction platform that directly links endogenous DNA repair activity to CRISPR-Cas12a activation. By coupling base excision repair (BER) events to a programmable DNA-based transducer, our system converts the activity of DNA glycosylases— such as uracil-DNA glycosylase (UDG) and human 8-oxoguanine glycosylase (hOGG1)—into a robust fluorescence signal via Cas12a-mediated *trans*-cleavage. This one-step CRISPR-based assay operates directly in cell lysates, enabling rapid and sensitive readout of enzymatic activity with high specificity. Additionally, it also enables in 15 minutes the throughput screening of novel potential inhibitors with high sensitivity. The modular design allows adaptation to diverse repair enzymes, offering a generalizable strategy for transforming intracellular repair events into programmable outputs. This approach lays the foundation for activity-based molecular diagnostics, synthetic gene circuits responsive to cellular states, and new tools for monitoring DNA repair in real time and drug screening.

## INTRODUCTION

Nature has evolved an extraordinary repertoire of reaction networks into interconnected communication pathways that efficiently catalyze multiple biochemical reactions in response to diverse stimuli and cellular conditions, ensuring proper metabolism and cellular reproduction.^1^ Recreating regulatory networks *in vitro* holds the potential to understand complex cellular processes and advance artificial technologies such as life-like materials,^2, 3^ artificial cells,^4, 5^ biosensing,^6-8^ and molecular computing.^9, 10^ To do so, precise spatiotemporal control of biomolecule interactions at the nano/micro-scale is necessary. In this respect, DNA nanotechnology offers unparalleled control over biomolecules by leveraging programmable nanoscale scaffolds^11-14^ to spatially organize antibodies,^15, 16^ enzymes^17-20^ and proteins.^21, 22^ In addition, dynamic chemical reaction networks based on synthetic DNA systems enable input-responsive tasks,^7, 23-27^ providing precise temporal control^28^ and regulations of downstream biological reactions.^29, 30^ Most artificial nucleic acid-based systems can be readily programmed using strand displacement reactions (SDRs). However, they still face notable limitations in integrating non-nucleic acid molecules and establishing efficient and sensitive *ligand-to-oligonucleotide* signal transduction networks.^31, 32^ While different transduction mechanisms - such as ligand-binding-induced proximity,^33-35^ structure switching^36^ and allostery^37, 38^ – that use synthetic DNA-based transducers (e.g., aptamers, protein-DNA conjugates, switches, etc.) have been developed, these systems generally depend on the inherent binding affinity between the ligand and the synthetic receptor. This limits signal amplification unless coupled with enzyme-free nucleic acid amplification strategies, which often require complex molecular design and exhibit unintended signal leakage.^39^

Advances in synthetic and molecular biology have spawned new tools, such as transcriptional^40, 41^ and CRISPR^42-44^ sensors, that integrate input-responsive DNA systems with downstream enzymatic amplification, enabling the transduction of biological inputs into amplified outputs. In particular, CRISPR-based platforms harness the collateral (*trans*-) cleavage activity of type V and VI CRISPR effector proteins (e.g. Cas12 and Cas13 enzymes) to achieve high sensitivity through signal amplification.^42, 45^ Among these, the Cas12a system is activated upon recognition of double and single-stranded DNA targets complementary to its CRISPR RNA (crRNA) guide, resulting in a fluorescence output via non-specific *trans-*cleavage activity of FRET-based DNA reporters.^46, 47^ Despite the versatility, these systems are limited by the strict requirement for sequence complementarity with the crRNA, which poses challenges for detecting non-nucleic acid targets.^48^ To address this limitation, conditional CRISPR-Cas systems regulated by external molecular inputs have been developed, offering promising strategies to partially overcome this limitation.^35, 49-52^ However, to date, direct regulation of CRISPR activity by upstream enzymatic activity has not yet been demonstrated.

Among potential upstream regulators, the DNA repair machinery - which restores chemically altered DNA (i.e., “DNA damage”) - represents a particularly promising system for triggering CRISPR-based responses. DNA repair enzymes act on DNA substrates, which can be rationally designed to become functional CRISPR targets only after enzymatic repair. This strategy offers a novel route to couple DNA repair activity with downstream CRISPR activation, enabling ultrasensitive and activity-dependent monitoring of DNA repair enzymes. To date, current methods for assessing repair capacity rely largely on indirect approaches, such as measuring gene or protein expression levels or single nucleotide polymorphism (SNP) screening. However, these do not always correlate with actual enzymatic activity, and SNPs can be uninformative when genes are not expressed.^53, 54^ Cell-based assays like the Comet assay^55^ or host cell reactivation^56^ (HCR) can assess repair capacity but often require complex experimental setups to isolate specific repair pathways in cells and tissues. Although DNA repair activity strongly influences the response to DNA-damaging chemotherapeutics,^57^ and repair inhibitors have been explored to boost drug efficacy,^58^ real-time, direct monitoring tools for specific repair enzymes are still lacking. Chemically-modified nucleic acid probes of DNA repair provide a promising alternative,^59^ allowing direct visualization of repair in cells^60, 61^ and compatibility with high-throughput screening of drugs in biological media. However, their clinical utility remains limited by challenges in probe design, synthesis, and relatively high detection thresholds due to the absence of signal amplification.^62^

Motivated by the above arguments, we report here on a strategy to *rewire* the activity of DNA repair enzymes – typically limited to correcting damaged bases – into a functional trigger for CRISPR-Cas12a signal transduction through a synthetic DNA-based molecular device (hereafter referred to as *DNA Transducer*, Figure 1). Our approach exploits the high specificity of DNA glycosylases (i.e., UDG and hOGG1), which recognize and excise rationally introduced lesions within the DNA Transducer. The repair activity induces a molecular reconfiguration of DNA Transducer that, in turn, activates Cas12a collateral cleavage only upon successful repair. By reprogramming DNA repair into a CRISPR-powered synthetic output in a one-step assay, we establish a one-pot, rapid detection platform capable of real-time monitoring of base excision repair activity directly in cell lysates. Furthermore, we demonstrate that our assay can be readily adapted for throughput screening of DNA repair enzyme inhibitors.

**Figure 1.**
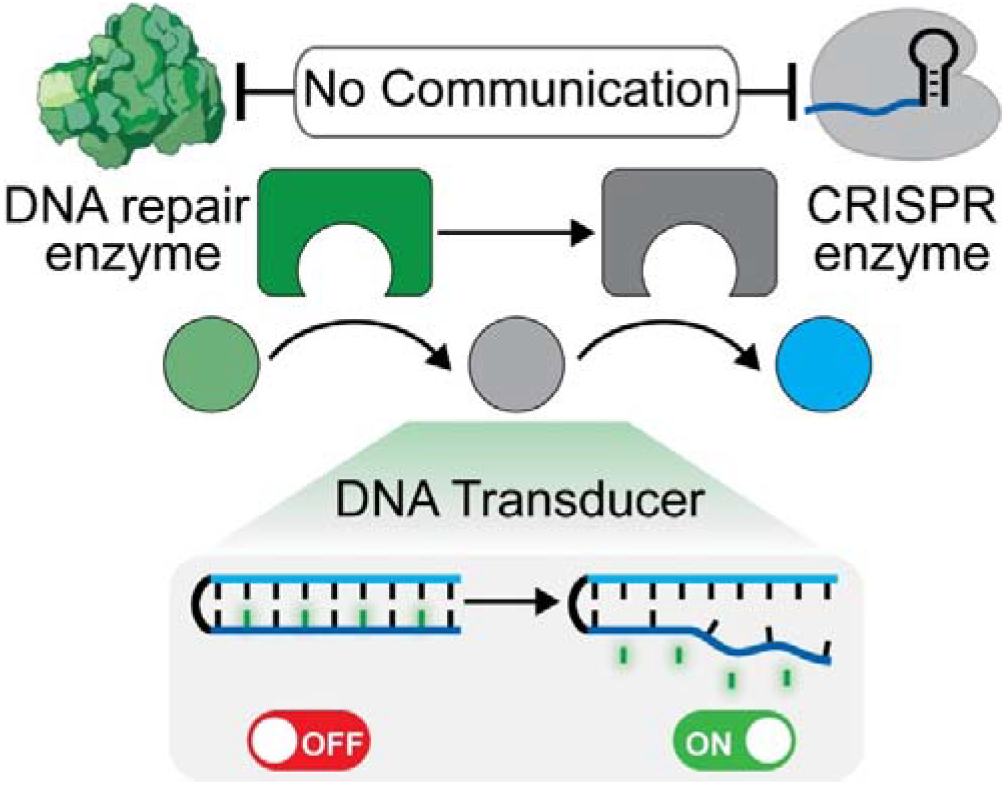
Schematic of artificial communication pathway between DNA repair and CRISPR-Cas12a enzyme controlled by a DNA-based molecular transducer. In its native hairpin state, the DNA Transducer is the substrate of a DNA repair enzyme as it contains one or more damaged nucleotides. Once a site-specific BER enzyme exerts its activity on the mutated base(s) of the DNA Transducer, forming abasic sites (or nicks, here not shown), the structural destabilization of the hairpin enables the binding of the RNP complex and Cas12a activation.

## RESULTS AND DISCUSSION

### Synthetic DNA Transducers enabling DNA repair-activated CRISPR signal transduction

We have rationally designed a set of synthetic DNA-based molecular transducers (DNA Transducer) that can be programmed to activate the CRISPR-Cas12a system only upon specific repair activity (Figure 1). Our DNA Transducer presents three specific features: i) it is a synthetic DNA hairpin designed to contain one or more *damaged* nucleotides (indicated by green symbols in Figure 1, e.g., Uracil or 8-oxoguanine, G-oxo) in its sequence; ii) it is recognized as a natural enzymatic substrate of the specific DNA repair enzyme; iii) it is rationally designed to populate two distinct conformational states, as it adopts a Cas12a-inactive hairpin structure (i.e. Cas12a-inactive state) that switches into an Cas12a-active conformation only upon repair activity. Specifically, our design takes advantage of the structural destabilization of the hairpin upon repair activity to effectively regulate Cas12a ribonucleoprotein (RNP) binding and activity. Notably, both our group and others have previously demonstrated that DNA hairpins containing Cas12a target sequences can be rationally engineered to resist Cas12a recognition and cleavage under native conditions.^35, 63^ Here, we designed a set of DNA Transducers containing base lesions into the non-target strand (NTS) embedded within the hairpin stem (dark blue sequence in Figure 1). These lesions are specifically targeted by the selected base excision repair (BER) enzyme, which recognizes the mutated bases and introduces abasic (AP) sites (UDG) or nicks (hOGG1), destabilizing the hairpin structure in response to glycosylase activity. Because of the specific repair activity - and consequent hairpin destabilization - the non-specific *trans-*cleavage activity of Cas12a is switched on, resulting in an amplified signal transduction due to the digestion of FRET-labelled DNA reporters.

### Design of UDG-responsive DNA Transducers downstream controlling Cas12a-based activity

As our test bed, we designed uracil DNA glycosylase (UDG)-responsive DNA Transducer (UDG-Transducers, Figure 2) downstream controlling CRISPR-Cas12a system. Uracil (U) is a common DNA damage introduced by dUMP misincorporation or cytosine deamination, resulting in U:A and U:G mispairs, respectively, which can compromise genomic stability,^64^ impacting human health.^65, 66^ UDG enzyme exhibits specific recognition and excision of uracil bases within DNA by hydrolyzing the N-glycosidic bond (Figure 2A, right panel), generating abasic sites that subsequently initiate the BER pathway. To detect UDG activity, we designed a set of UDG Transducers presenting a 20-nt long crRNA targeting portion (blue region, Figure 2B) with different numbers of uracil lesions (green symbols, from 0 to 4 U) in the non-target strand (NTS), embedded within the hairpin stem (dark blue motifs, Figure 2B).

**Figure 2.**
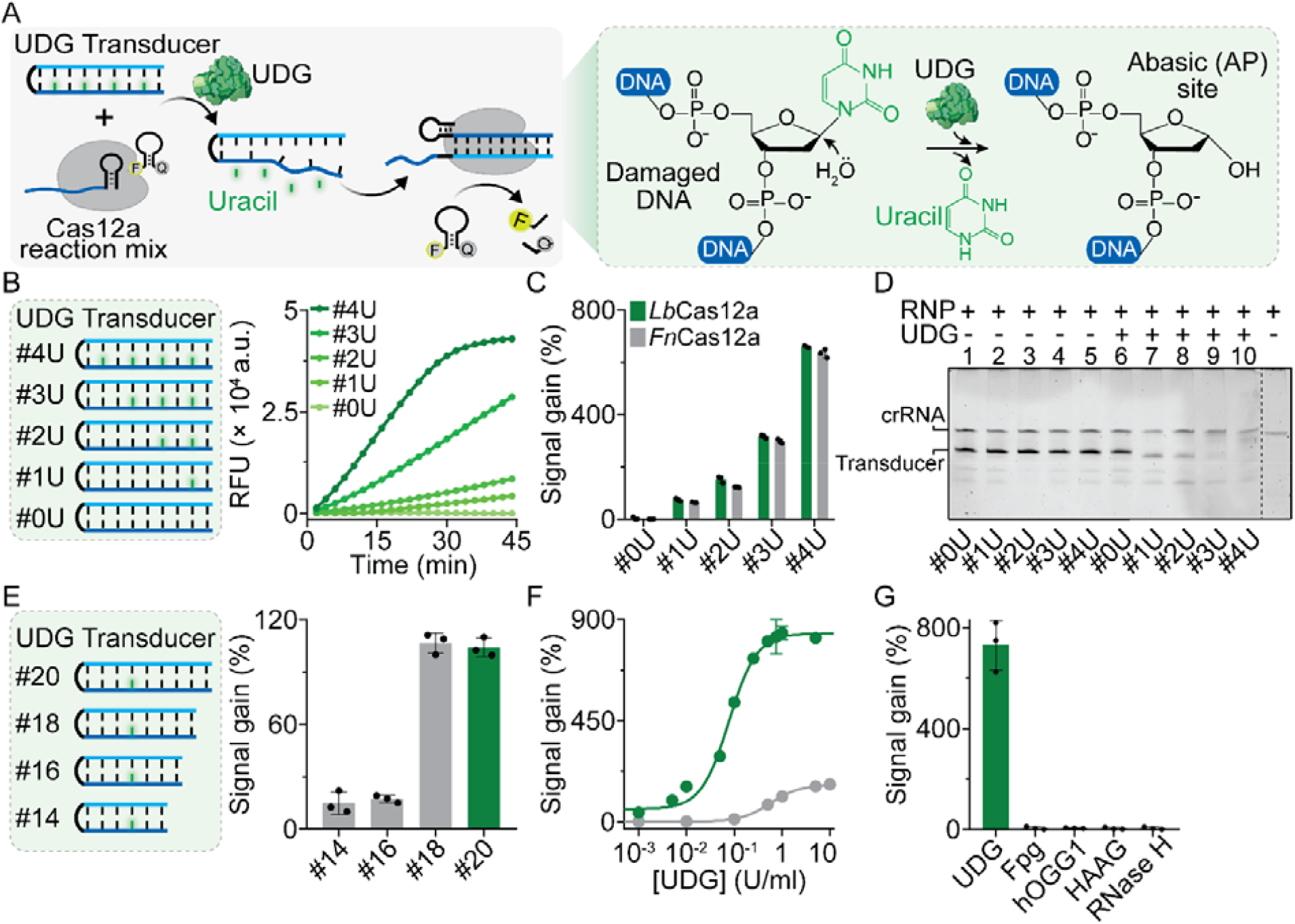
Design and optimization of UDG Transducer triggering Cas12a *trans*-cleavage activity upon hairpin reconfiguration induced by UDG activity. **A**) Schematic illustration of our sensing strategy for UDG activity monitoring. The mechanism of uracil excision by UDG is indicated on the right side. **B**) Fluorescence kinetic analysis over time showing different profiles of *Lb*Cas12a collateral cleavage activity triggered by different DNA Transducers (from #0U to #4U) that were pre-incubated with UDG (1 U/ml). The kinetic profiles indicate that the activation of *Lb*Cas12a depends on the number of uracil bases within the DNA Transducer (1 nM). **C**) Fluorescence signal gain generated by the addition of UDG (1 U/ml) in solution containing the DNA Transducer #4U (1 nM) using different Cas12a orthologs reaction mix. D) Denaturing PAGE analysis reporting the *cis*-cleavage activity of *Lb*Cas12a on UDG Transducers in the absence and presence of UDG (1 U/ml). **E**) Fluorescence signal gain obtained upon addition of 0.5 U/ml UDG (t = 15 min) using UDG Transducers with varying lengths of crRNA targeting region (from 14 to 20 nt). **F**) Dose-response curves of the *Lb*Cas12a-based UDG activity assay (green curve) compared to a direct fluorescent assay (gray curve) using a FRET-labelled UDG Transducer (500 nM). **G**) Specificity test at saturated concentrations of UDG (0.5 U/ml), and 10 U/ml concentration of non-specific target enzymes. Experiments were conducted at 37 °C by adding the Cas12a reaction mix (500 nM of FRET-based DNA reporter and 20 nM of Cas12a/crRNA complex) to a 25 μl buffer solution (10 mM Tris-HCl, 50 mM NaCl, 10 mM MgCl□, and 0.1 mg/ml BSA, at pH 7.9) containing UDG Transducer (1 nM) and a specific concentration of UDG. Signal gain (%) was calculated after 15 min of cleavage reaction, representing the relative fluorescence signal change associated with the Cas12a *trans*-cleavage activity upon UDG addition and relative to the background fluorescence obtained in the absence of UDG. Error bars represent the deviation from three independent experiments.

To study the Cas12a signal transduction activated by UDG activity, we performed collateral cleavage fluorescence assays by pre-incubation of UDG enzyme (1 U/ml) with different UDG Transducers (1 nM) and subsequent addition of a Cas12a reaction mix (500 nM of FRET-based DNA reporter and 20 nM of Cas12a/crRNA complex). As shown in Figure 2B, the UDG transducers containing uracil lesions function as regulatory switches for Cas12a, exhibiting increased fluorescence over time upon UDG addition (1 U/ml). As expected, the transducer without uracil in the sequence (UDG Transducer #0U) displays no fluorescence change. This confirms that the excision of U due to UDG activity induces hairpin destabilization, which results in partial or complete unfolding of the hairpin, enabling Cas12a binding and activation. Additionally, increasing the amount of uracil in the hairpin from 1 to 4 results in a faster Cas12a-based signal transduction. This is coherent with the fact that increasing the number of abasic sites (from 1 to 4) in the probe results in a more destabilized hairpin structure upon UDG repair activity, which facilitates Cas12a RNP binding and activation. These findings indicate that by simply controlling the number of uracil nucleotides, it is possible to modulate the thermodynamics of UDG Transducer unfolding, providing also an effective strategy for regulating Cas12a activity. To demonstrate the generalizability of our approach, we also tested different Cas12a orthologs from *Lachnospiraceae bacterium* (*Lb*Cas12a) and *Francisella novicida* (*Fn*Cas12a). Both orthologs show a similar fluorescence signal gain due to UDG activity, which is dependent on the number of U within the hairpin stem (Figure 2C). Next, we also studied site-specific (*cis*-)^67^ cleavage activity of Cas12a on UDG Transducers using denaturing PAGE (Figure 2D). Our results confirm that *cis*-cleavage activity is activated only by UDG Transducers containing uracil residues (UDG Transducer #1U to #4U) pre-incubated with UDG (lanes 7 to 10, Figure 2D). To confirm the proposed mechanism, we performed an activity-based assay using the same DNA Transducer containing four uracil lesions that was terminally labeled with FAM and BHQ1 (UDG Transducer #4U-FQ, 500 nM) to generate a direct fluorescence signal transduction. As expected, the addition of UDG (10 U/ml) results in an increase of fluorescence induced by uracil excision and hairpin structure destabilization, whereas the same DNA Transducer containing four AP sites (UDG Transducer #4AP-FQ) does not show any signal change over time in the presence of UDG (Figure S1). Melting denaturation assays also confirm that UDG activity leads to destabilization of hairpin structure of the UDG Transducer #4U (from Tm = ∼ 85 °C in the absence of UDG to T_m,_ = 59.6 °C upon UDG activity, Figure S2).

As Cas12a collateral activity depends on the length of the double stranded DNA targeting region, we also tested a set of UDG Transducers with crRNA-targeting lengths ranging from 14 to 20 nucleotides, each containing a single uracil lesion (Figure 2E). The observed signal gain induced by UDG activity (1 U/ml) aligns with previous studies indicating that optimal collateral cleavage activity is achieved using 20-bp long DNA targeting motifs.^46^ Once identified the UDG Transducer #4U as optimal probe for rapid and sensitive Cas12a signal transduction, we further optimized its concentration to improve signal to noise ratio, as its concentration may affect the background signal, which in turn determines the overall signal change (1 nM, Figure S3). Under optimized experimental conditions, we performed Cas12a-based UDG activity assays to determine the limit of detection (LOD) of our platform (0.0009 U/ml, RSD of 3.4 %, Figure 2F). Comparing the proposed detection system with a direct activity-based FRET assay using the same transducer terminally labelled with a FRET pair (DNA Transducer #4U-FQ, 500 nM), we highlight that our Cas12a-based detection system results in a 49.8-fold increase in sensitivity (LOD_#4U-FQ_ = 0.02 U/ml). As expected, the platform exhibits high specificity, as no significant signal variation is observed in the presence of non-specific DNA repair enzymes (Figure 2G). Finally, we also tested the method against human single-strand selective monofunctional uracil-DNA glycosylase (hSMUG1), further validating the versatility and robustness of our method (Figure S4).

### Cas12a-based real-time monitoring of UDG activity in human cell lysate

Motivated by the above results, we tested our Cas12a-based UDG activity assay in HEK293T cell lysate (Figure 3A). First, we tested UDG-inactive HEK293T cell lysate by adding Cas12a reaction mix in the solution containg the DNA Transducer #4U and spiked UDG. Our assay can detect UDG activity in HEK293T cell lysate in the dynamic range from 0.0085 to 0.65 U/ml (K_1/2_ = 0.07 ± 0.01 U/mL, Figure 3B) after 15 minutes of reaction (Figure S5). To evaluate the accuracy of the Cas12a-based method, we conducted recovery tests by spiking UDG at four different concentrations into HEK293T cell lysate, confirming that our approach can be used for detecting its activity in complex biological specimens (Figure 3C). Then, to demonstrate real-time monitoring of the UDG activity, we used HEK293T lysate with overexpressed uracil-N-glycosylase (UNG) activity. To do so, subsequent dilution of HEK293T lysate was tested by simply adding Cas12a reaction mix with the UDG Transducer #4U, confirming highly sensitive detection (Figure 3D). To further confirm that the Cas12a reaction network is exclusively triggered by UNG activity in the cell lysate, we conducted a control experiment in the presence of the uracil-DNA glycosylase inhibitor (UGI). UGI specifically inhibits UNG by forming a stable, irreversible UNG:UGI protein complex in a 1:1 stoichiometry, thereby preventing UNG from excising uracil from DNA.^68^ As expected, by increasing the concentrations of UGI in the cell lysate (10 μg/ml), a dose-dependent reduction in fluorescence intensity is observed (Figure 3E). These results indicate the capacity of the assay to enable either real-time monitoring of glycosylase activity and also inhibitor screening directly in cell lysate.

**Figure 3.**
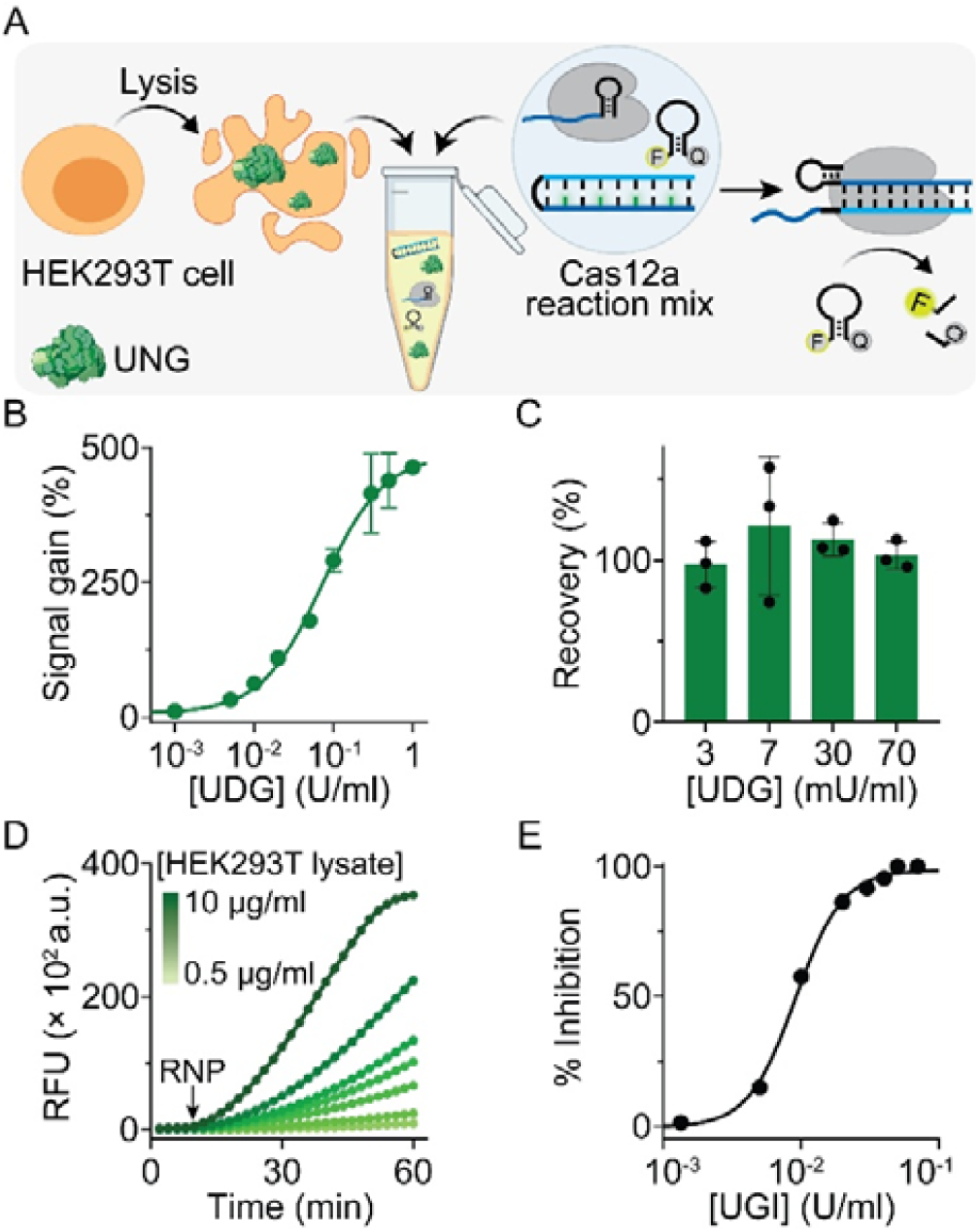
Cas12-based UDG/UNG activity assay in human HEK293T lysate. **A**) Schematic illustration of the Cas12a-based reaction network for UNG activity monitoring in human HEK293T cells. **B**) Dose-response calibration curve of UDG activity in human HEK293T cells obtained using the optimized Cas12a-based reaction mix (DNA Transducer #4U 1 nM, Cas12a-crRNA 20 nM and FRET-based DNA reporter 500 nM). **C**) Recovery test in HEK293T cell lysate achieved by spiking with various concentrations of UDG. **D**) Fluorescence kinetic traces of Cas12a collateral activity in human HEK293T cell lysate overexpressing UNG. **E**) Inhibition plot of UNG activity using UNG-overexpressing HEK293T cell lysates and the Cas12a reaction mix (DNA Transducer #4U 1 nM, Cas12a-crRNA 20 nM and FRET-based DNA reporter 500 nM) in the presence of varying concentrations of UGI inhibitor. Recovery tests and inhibition experiments were conducted according to the experimental procedures reported in the Supporting Information. Error bars represent the deviation from three independent experiments.

### Synthetic DNA Transducers rewire hOGG1 activity into Cas12a Signal Amplification

Our strategy is highly versatile, as it enables monitoring of various classes of glycosylases by simply changing the specific DNA damage within the DNA Transducer. As a proof-of-concept demonstration, we tested human 8-oxoguanine DNA glycosylase 1 (hOGG1) which a bifunctional glycosylase that not only excises damaged bases but also successively cleave phosphodiester bonds at the generated abasic sites.^69^ hOGG1 is a key enzyme involved in repairing oxidative DNA damage that specifically detects 8-oxoguanine (8-oxoG), an oxidative DNA lesion that can cause G:C→A:T transversion mutations if unrepaired.^70, 71^ Given the central role of oxidative DNA damage in the development of cancer^72, 73^ neurodegenerative diseases^74^, and aging^75^, monitoring OGG1 activity could provide valuable insights into these conditions and their progression. Here, we designed a hOGG1 Transducer having the same Cas12a-targeting region used for UDG monitoring but containing a single 8-oxoG lesion in the middle of the crRNA targeting region (red symbol, Figure 4A). This design makes it possible to use the same Cas12a reaction mix for detecting both UDGs and hOGG1, thereby confirming the generalizability of the sensing approach. Specifically, when hOGG1 excises 8-oxoG from the DNA molecular transducer (hOGG1 Transducer), the secondary lyase activity results in the nicking of the substrate, leading to a *release* of the short DNA motif (9 nt) from the hairpin structure. This promotes overall hairpin destabilization, which facilitates Cas12a RNP binding to the unpaired nucleotides of the target strand of the hOGG1 Transducer, ultimately activating Cas12a *trans*-cleavage activity and degradation of the fluorescence reporter (Figure 4A).

**Figure 4.**
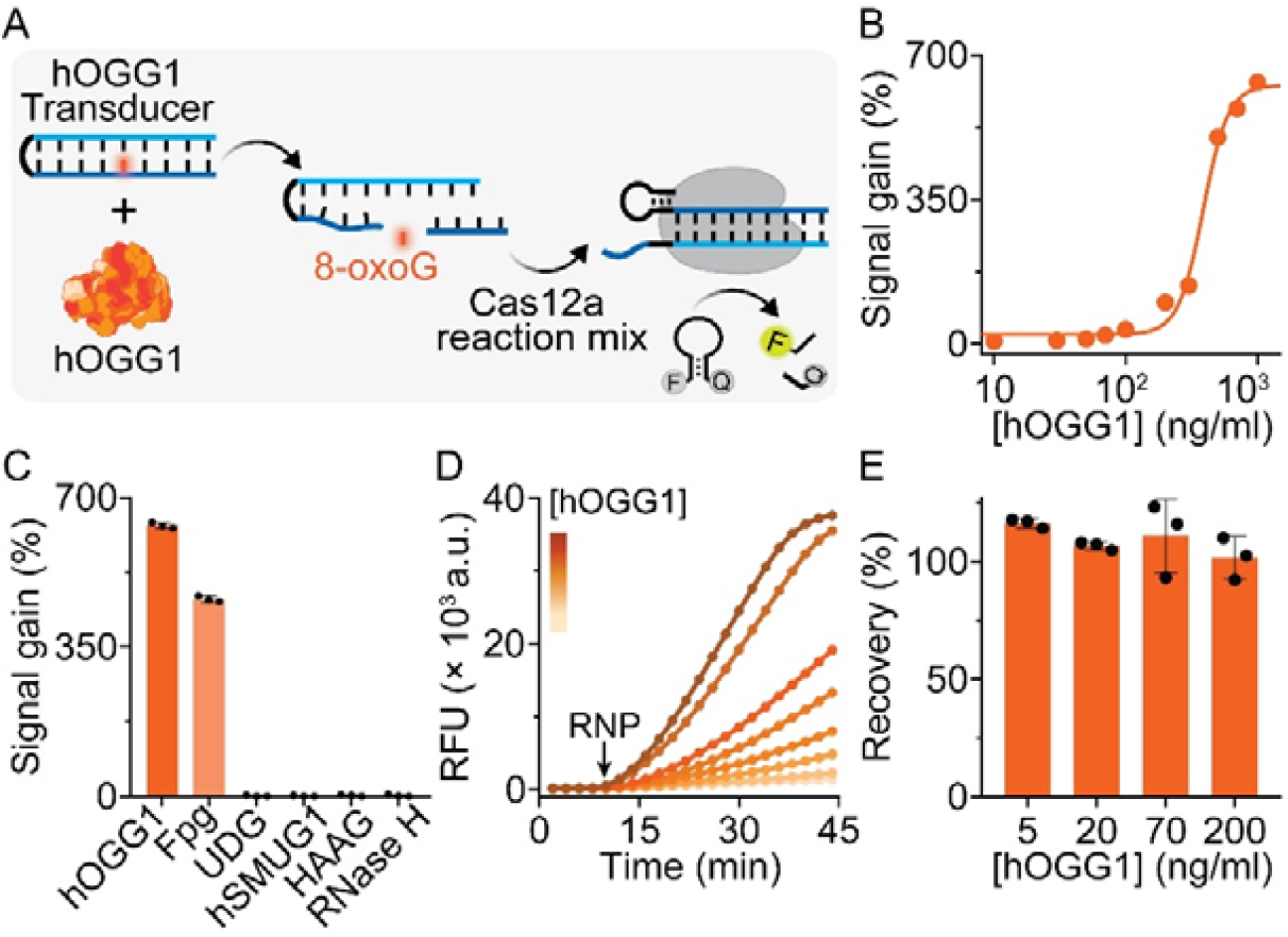
Cas12a-powered activity assay for the monitoring of hOGG1 activity. **A**) Scheme of our sensing strategy based on an 8-oxoG-containing DNA Transducer (hOGG1 Transducer) that triggers Cas12a-based signal transduction upon repair activity. **B**) Calibration curve obtained using the hOGG1 Transducer (1 nM) by adding increasing concentrations of hOGG1 and the Cas12 reaction mix. **C**) Specificity test in the presence of non-specific enzymes (10 U/ml). **D**) Fluorescence kinetic traces of Cas12a activation in HEK293T cell lysate supplemented with increasing concentrations (3, 10, 30, 50, 100, 150, 300 and 500 ng/ml) of hOGG1. **E**) Recovery test in HEK293T cell lysate spiked with different concentrations of hOGG1. Recovery tests were conducted according to the experimental procedures reported in the Supporting Information. Error bars represent the deviation from three independent experiments.

We first characterized our assay using E. coli formamidopyrimidine-DNA glycosylase (Fpg), a bacterial analogue of hOGG1.^76^ Also in this case, denaturing PAGE confirms that the *cis*-cleavage activity of Cas12a is present only on the DNA Transducer containing the 8-oxoG lesion that is repaired by the Fpg activity (lane 4, Figure S6). In contrast, the DNA Transducer lacking the 8-oxoG lesion (i.e. hOGG1 Transducer w/o Goxo) remains undigested even in the presence of Fpg (Lanes 5-8). Our Cas12a-based Fpg activity assay exhibits high sensitivity with a limit of detection (LOD) of 0.001 U/ml for Fpg, outperforming direct activity-based assays (LOD = 0.3 U/ml) under the same conditions (Figure S7). We then applied our CRISPR platform to detect hOGG1, confirming sensitive detection (LOD = 14.2 ng/ml, Figure 4B) and high specificity (Figure 4C). Our CRISPR-based platform was further assessed in human HEK293T cell lysates through spike and recovery assays by adding known concentrations of hOGG1 (Figure 4D), following dose-response fluorescence kinetics over time. Statistical analysis revealed a linear detection range of 3 to 300 ng/mL, with an R^2^ value of 0.995 (Figure S8). Additionally, we assessed the recovery percentages of spiked hOGG1 at four different concentrations, achieving recovery rates ranging from 101 % to 115 % (Figure 4E), confirming the accuracy of our single-step detection method.

### Cas12a-based throughput screening of the hOGG1 inhibitors

Recent studies indicated that the inhibition of hOGG1 is a promising strategy for developing treatments for cancer and inflammation.^77, 78^ Leveraging the high sensitivity and rapid response of our single-step CRISPR-based hOGG1 activity assay, we re-adapted our system for throughput screening of small molecule inhibitors (Figure 5A). To this end, we tested several well-characterized small-molecule inhibitors targeting hOGG1 and Fpg. Specifically, we assessed two inhibitors of hOGG1 operating through different mechanisms. Indeed, O8 (Inh#1) prevents catalytic AP lyase activity of hOGG1 without affecting protein-substrate binding,^79^ while SU0268 (Inh#2) inhibits both DNA binding and base excision activity of the enzyme.^80^ Furthermore, we examined three 2-thioxanthine (2TX) derivatives (Inh#3, #4 and #5), known to irreversibly inhibit zinc finger (ZnF)-containing Fpg/Nei DNA glycosylases.^81, 82^ We tested a range of inhibitor concentrations (0.1 μM to 350 μM) to assess the inhibition of hOGG1 and Fpg. Consistent with previous studies,^79, 80^ hOGG1 inhibition exhibits a selective dependence on the concentrations of Inh#1 (O8) and Inh#2 (SU0268), yielding IC50 values of 2.5 ± 0.5 μM and 28.6 ± 7.3 μM, respectively (Figure 5B). In contrast, Inh#3 and Inh#5 selectively inhibit Fpg (Figure S14). On the other hand, Inh#4 shows dual inhibition, affecting both hOGG1 (IC50 = 11.9 ± 0.4 μM) and Fpg (IC50 = 70.3 ± 0.7 μM) (Figure 5B and S9). The heatmap reported in Figure 5C offers a concise overview of the inhibition effects at a fixed concentration of 200 μM, highlighting the different activity between the bacterial and human enzymes, and confirms the selective inhibition profiles of each inhibitor. Additionally, it confirms that our Cas12a-based screening can be used to distinguish between various inhibitors based on their target specificity. As a control, we confirmed that Cas12a activity is not affected by the presence of 10% DMSO - that is required for inhibitor solubilization – and the Cas12a signal off is due exclusively to the glycosylase inhibition (Figure S10 and S11). It is important to highlight that the single-step CRISPR-based screening is performed in a 96-well plate microvolume (25 ul) and takes approximately 15 min after the addition of RNP. It costs approximately less than 0.7 euro per test as it requires a minimal amount of DNA Transducer (1 nM) and Cas12a reaction mix (20 nM RNP and 500 nM FRET-based DNA reporter), thanks to the sensitive nature of the assay.

**Figure 5.**
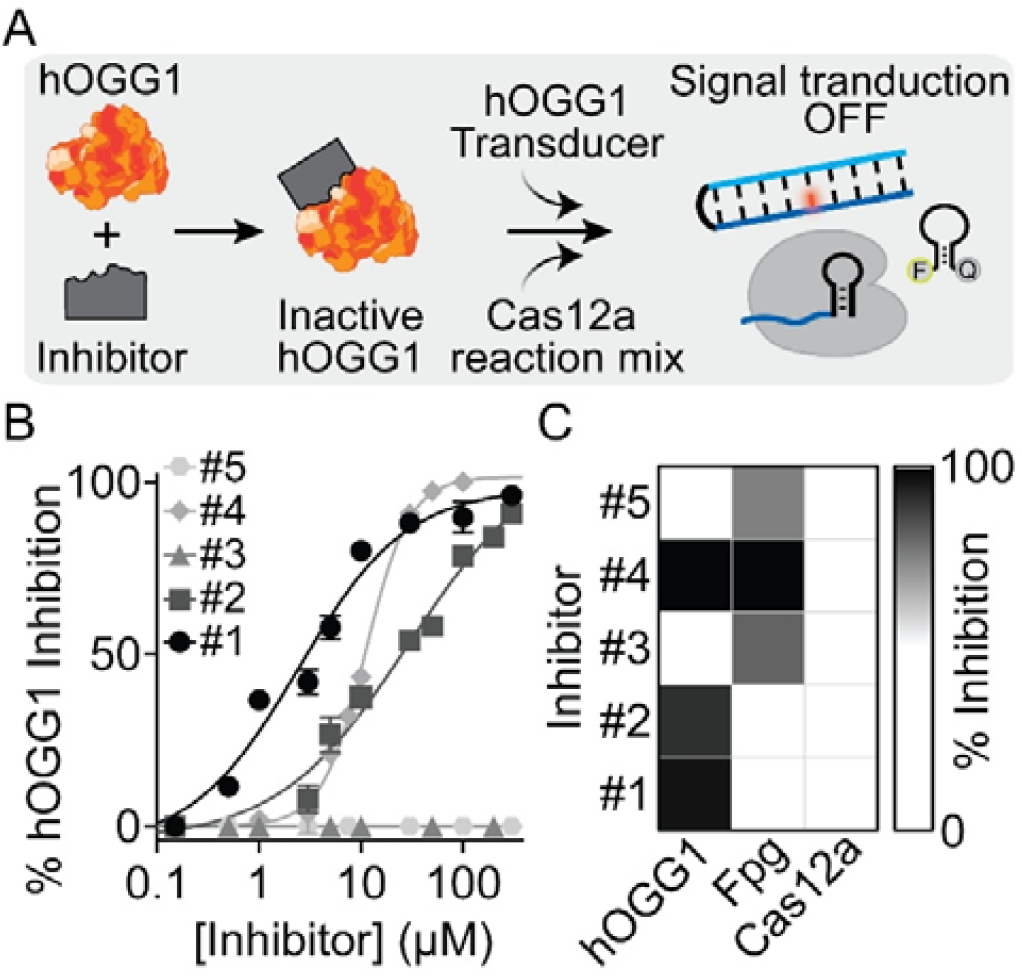
Cas12a-based throughput screening of hOGG1 inhibitors. **A**) Schematic description of the assay. **B**) Plot of % inhibition of hOGG1 activity at various concentrations of different inhibitors. **C**) Heatmap showing % inhibition observed with different inhibitors (200 μM) on hOGG1, Fpg and Cas12a/crRNA complex (20 nM), as a control experiment. In (B) and (C), Inh#1 to #5 represent O8, SU0268, 2-thioxanthine, Mercapto-6,8-purinediol, and 8-mercaptoadenine, respectively. Experiments were conducted at 37 °C by adding the Cas12a reaction mix (500 nM of FRET-based DNA reporter and 20 nM of Cas12a/crRNA complex) to a 25 μl buffer solution (10 mM Tris-HCl, 50 mM NaCl, 10 mM MgCl□, and 0.1 mg/ml BSA, at pH 7.9) containing hOGG1 Transducer (1 nM), hOGG1 (700 ng/ml) and varying concentration of inhibitors. Inhibition (%) for each inhibitor was calculated after a 15-minute cleavage reaction, as described in the Supporting Information. Error bars represent the standard deviation from three independent experiments.

Overall, these findings underscore the potential of our CRISPR/Cas12a-based throughput screening for its adaptation in high-throughput automatic format, showcasing potential also for selective inhibitor identification and broader applications in DNA repair-targeted therapies.

## Conclusions

In this study, we developed synthetic DNA Transducers that activate CRISPR-Cas12a upon enzymatic repair, enabling rapid, real-time monitoring of DNA glycosylase activity. By harnessing the substrate specificity of repair enzymes for damaged DNA bases, we show that their activity can trigger Cas12a-mediated signal transduction, effectively converting DNA damage recognition into amplified fluorescence output. This one-step assay couples lesion excision with Cas12a *trans*-cleavage to achieve ultrasensitive detection of glycosylase activity directly in complex biological samples such as cell lysates.

Our platform overcomes several limitations of conventional activity-based assays, including the need for chemically modified (unnatural) DNA probes, low sensitivity, and multistep workflows. Unlike previous CRISPR-based strategies, our system requires no auxiliary enzymes, complex probe architectures, or multi-reagent additions as an amplified output is generated by simply mixing components in solution. The assay is built on DNA hairpins that serve as natural substrates for the targeted repair enzymes, and its simple design allows easy adaptation to different DNA lesions (e.g., uracil or 8-oxoG), enabling broad applicability across the DNA repair landscape. Our assay is indeed designed to generate an output by simply mixing the reagents in solution without complex, multistep and reagent-intensive procedures. Beyond its sensing capabilities, this platform offers a powerful tool for throughput screening of DNA repair inhibitors. We demonstrate this by evaluating several small-molecule inhibitors of hOGG1 and Fpg. Ultimately, our scalable and efficient approach not only facilitates fundamental studies of DNA repair mechanisms but also supports the development of targeted therapeutics, real-time biosensors, and drug discovery tools.

## AUTHOR INFORMATION

### Author Contributions

All authors have given approval to the final version of the manuscript. N.B. conceived, designed, and performed experimental work, analyzed data, and wrote the first draft. A.B. and R.M. discussed the A.P.

### Notes

The authors declare no competing financial interest.

## ACKNOWLEDGMENT

The research leading to these results has received funding from AIRC under MFAG 2022 - ID. 27151 project – P.I. Porchetta Alessandro. A.P. acknowledge financial support under the National Recovery and Resilience Plan (NRRP), Mission 4, Component 2, Investment 1.1, Call for tender No. 104 published on 2.2.2022 by the Italian Ministry of University and Research (MUR), funded by the European Union – NextGenerationEU– Project Title “CRISPR-Cas-based sensing platforms for the monitoring of clinically relevant antibodies”– CUP D53D23009090001-Project Code 2022FPYZ2N - Grant Assignment Decree No. 958 adopted on 30-06-2023 by the Italian Ministry of Ministry of University and Research (MUR) and by “PNRR M4C2-Investimento 1.4-CN00000041” financed by NextGenerationEU. N.B. was supported by a Fondazione Umberto Veronesi postdoctoral fellowship.

