## Supplementary material for "Rewiring DNA Repair Activity into CRISPR Signal Transduction via Synthetic DNA Transducers": SI_03.07.2025.docx

#### Experimental Procedures

#### Reagents and Materials

Sodium chloride (NaCl), sodium hydroxide (NaOH), magnesium chloride (MgCl_2_), Trizma hydrochloride (Tris–HCl), Trizma, boric acid, ethylenediamine tetraacetic acid (EDTA), acrylamide/bis-acrylamide 40% solution, ethylenediamine tetraacetic acid (EDTA), N,N,N′,N′-tetramethyl ethylenediamine (TEMED), ammonium persulfate (APS), DEPC-treated water, bovine serum albumin (BSA), 2-mercapto-6,8-purinediol, 8-mercaptoadenine, Dimethyl Sulfoxide (DMSO). 2-thioxanthine and Gel Loading Buffer II were purchased from ThrmoFisher Scientific. Invitrogen provided SYBR Gold nucleic acid stain. OGG1 inhibitors O8 and SU0268 were purchased from MedChemExpress. Uracil DNA glycosylase (UDG), uracil glycosylase inhibitor (UGI), E. coli formamidopyrimidine DNA glycosylase (Fpg), human single-strand selective monofunctional uracil-DNA glycosylase 1 (hSMUG1) and EnGen® Lba Cas12a from *Lachnospiraceae* bacterium ND2006 were purchased from New England Biolabs (Beverly, MA, USA). FnCas12a from *Francisella novocida* bacterium was generously provided by the laboratory of Professor Giuseppe Perugino at the University of Naples "Federico II" which was produced as previously reported in our work.^1^ Human 8-oxoguanine DNA glycosylase (OGG1), transient overexpression lysate of uracil-DNA glycosylase (UNG), and HEK293T cell lysate were purchased from Origene. The enzymes were aliquoted and stored at -20 °C or -80 °C.

#### Oligonucleotides

All the oligonucleotides employed in this work were synthesized, labeled, and HPLC-purified by Metabion International AG (Planegg, Germany) and used without further purification. DNA and RNA oligonucleotides were dissolved to 100 μM in 100 mM Tris-Hcl pH 7.5, and DEPC-treated water, respectively. All oligonucleotides were aliquoted and stored at -20 °C until use. The final concentration of the oligonucleotides was confirmed using Thermo Scientific™ Multiskan SkyHigh Microplate Spectrophotometer.

#### Fluorescence Experiments

Fluorescence kinetic measurements were performed in a 25 µl buffer solution containing 10 mM Tris-HCl, 50 mM NaCl, 10 mM MgCl₂, and 0.1 mg/ml BSA, at pH 7.9 unless otherwise specified. The measurements were carried out using a Stratagene Mx3005P Real-Time PCR System (Agilent Technologies, Santa Clara, CA) with 96-well PCR plates and optically transparent caps (Thermo Scientific). The fluorescence signals were recorded every 2 minutes, starting from the initial 2-minute time point, at a constant temperature of 37 °C using a filter set optimized for FAM detection.

For fluorescence melting assays, a 1 ml cuvette and a JASCO PF-8050 spectrofluorometer with excitation wavelength set at 490 (±5) nm and acquisition at 520 (±5) nm were used. Detailed procedures employed in the different experiments are reported below.

#### Fluorescence *Trans*-Cleavage Assays

Fluorescence *trans*-cleavage experiments shown in Figure 2 were conducted by adding a concentration of UDG to the buffer solution (10 mM Tris-HCl, 50 mM NaCl, 10 mM MgCl_2_ 0.1 mg/ml BSA, pH 7.9) containing 1 nM UDG DNA Transducer (see Table S1) and 500 nM FRET-based DNA reporter. An aliquot of 22.5 µl of this mixture was transferred to the 96-well PCR plate, followed by the addition of 2.5 µl of the Cas12a/crRNA (RNP) complex, which had been pre-incubated at 37 °C for 30 minutes with 200 nM of both Cas12a and crRNA, achieving a final concentration of 20 nM. Fluorescence intensity was then immediately monitored over time.

For the experiments shown in Figure 4, target enzyme (hOGG1 or Fpg) to a solution containing 1 nM hOGG1 DNA Transducer (see Table S2) and 500 nM FRET-based DNA reporter. After transferring 22.5 µl of the sample to a 96-well PCR plate and collecting fluorescence for 10 minutes, the RNP complex was added at a final concentration of 20 nM, and fluorescence intensity was immediately monitored over time. Inhibition assays shown in Figure 5 involved incubating inhibitors with 700 ng/ml hOGG1 or 0.75 U/ml Fpg in the buffer solution containing 10 mM Tris-HCl, 50 mM NaCl, 10 mM MgCl_2_ 0.1 mg/ml BSA, 10% DMSO, pH 7.9. After 30 minutes of incubation at 37 °C, 1 nM hOGG1 DNA Transducer (see Table S2) and 500 nM FRET-based DNA reporter were added to the reaction mixture. An aliquot (22.5 µl) of the sample was then transferred to a 96-well PCR plate and fluorescence was collected for 10 minutes. The RNP complex was then added at a final concentration of 20 nM, and fluorescence intensity was immediately monitored over time.

#### Cell lysate analysis

Experiments shown in Figures 3B and 3C were conducted using 10 µg/ml HEK293T cell lysate in a buffer solution (10 mM Tris-HCl, 50 mM NaCl, 10 mM MgCl_2_ 0.1 mg/ml BSA, pH 7.9). Different concentrations of UDG were spiked into the lysate sample, along with 1 nM UDG DNA Transducer (#4U) and 500 nM FRET-based DNA reporter. After transferring 22.5 µl of the sample to a 96-well PCR plate, fluorescence was collected for 10 minutes. The RNP complex was then added to a final concentration of 20 nM, and fluorescence intensity was immediately monitored over time. To assess measurement accuracy, recovery experiments were performed by spiking known concentrations of UDG (e.g., 0.003, 0.007, 0.03, and 0.07 U/ml) into the lysate sample. The recovered concentrations were determined using linear regression and compared with the added concentrations. Similar experiments were conducted by spiking different concentrations (e.g., 5, 20, 70, 200 ng/ml) of hOGG1 to the lysate sample along with the 1 nM hOGG1 DNA Transducer and 500 nM FRET-based DNA reporter (Figure 4D and 4E).

For the experiments shown in Figure 3D, the transient overexpression lysate of UNG was diluted to various concentrations in the buffer solution. To these lysate samples, 1 nM UDG DNA Transducer (#4U) and 500 nM FRET-based DNA reporter were added. Without any incubation, an aliquot of 22.5 µl of the mixture was transferred directly to a 96-well PCR plate, and fluorescence was collected for 10 minutes. The RNP complex was then added to achieve a final concentration of 20 nM, and fluorescence intensity was immediately monitored over time.

To test UGI inhibitor activity shown in Figure 3E, different concentrations of UGI were spiked into the transient overexpression lysate of UNG (10 µg/ml). After 40 minutes of incubation at 37 °C, 1 nM UDG DNA Transducer (#4U) and 500 nM FRET-based DNA reporter were added to the reaction mixture. An aliquot (22.5 µl) of the sample was then transferred to a 96-well PCR plate and fluorescence was collected for 10 minutes. The RNP complex was then added at a final concentration of 20 nM, and fluorescence intensity was immediately monitored over time.

#### Conventional FRET Assays

Conventional fluorescence assays were conducted by adding varying concentrations of UDG or Fpg to a buffer solution (10 mM Tris-HCl, 50 mM NaCl, 10 mM MgCl₂, 0.1 mg/ml BSA, pH 7.9) containing 500 nM FRET-labeled UDG DNA Transducer (#4U-FQ) or FRET-labeled hOGG1 DNA Transducer, respectively. An aliquot of 25 µl of the mixture was then transferred to a 96-well PCR plate, and fluorescence intensity was monitored over time.

#### Denaturing PAGE

*Cis*-cleavage assays shown in Figure 3D were performed by incubating different UDG DNA Transducers (see Table S1) and 50 nM of RNP complex in buffer solution (10 mM Tris-HCl, 50 mM NaCl, 10 mM MgCl_2_, 0.1 mg/ml BSA, pH 7.9) in the absence (lane 1 to 5) and in the presence (lane 6 to 10) of 1 U/ml UDG. To each 10 µl sample, 1 µl of Gel Loading Buffer II and 5 µl of formamide were added, and the samples were then boiled for 5 min. The samples were loaded onto a 20% denaturing polyacrylamide gel, which was run at room temperature in 1× TBE buffer (pH 8) at 120 V. The experiments were conducted using a Mini-PROTEAN Tetra cell electrophoresis unit (Bio-Rad) with a Bio-Rad PowerPac Basic power supply. The gel was stained with SYBR Gold and imaged using a ChemiDoc Imaging System (Bio-Rad) to visualize the DNA bands.

#### Melting assays

Fluorescence melting curves (Figure S2) were obtained by heating buffer solution (10 mM Tris-HCl, 50 mM NaCl, 10 mM MgCl₂, 0.1 mg/ml BSA, pH 7.9) containing 500 nM FRET-labeled UDG DNA Transducers, #4U-FQ and #4AP-FQ, either in the presence or absence of UDG, from 20°C to 90°C at a rate of 1°C/min. Fluorescence was monitored at 0.5 °C intervals using excitation at λₓ = 488 nm and emission at λₑₘ = 520 nm. Before the melting analysis, the solutions were annealed by heating to 90°C and then slowly cooling to room temperature. The reported melting curves were normalized using an interpolation model to estimate the melting temperature (Tₘ) for each experiment.

#### Data Analysis

Signal gain (%) is calculated at 15 minutes from the addition of the RNP complex to the reaction mixture unless otherwise stated. It represents the relative fluorescence signal change upon the addition of the target, relative to the background fluorescence obtained in the presence of the DNA Transducer. The signal gain is calculated using the following formula:

$$Signal gain \left( \% \right)= \frac{F_{target}- F_{0}}{F_{0}} \times100$$

where F_target_ is the fluorescence signal measured upon addition of the target enzyme and F_0_ is the fluorescence signal generated by the DNA Transducer in the absence of the target enzyme, respectively. Plots of signal gain (%) vs concentration of target enzyme were fitted with the following four parameters logistic equation:

$$Signal gain \left( \% \right)= B_{min} + \left( B_{max} - B_{min} \right)\frac{\left[ Target enzyme \right]^{n_{H}}}{K_{\frac{1}{2}}^{n_{H}} + \left[ Target enzyme \right]^{n_{H}}}$$

Where B_min_ and B_max_ are the minimum and maximum signal gain (%) values, respectively; K_1/2_ is the concentration of the target at a half-maximum signal gain (%), [Target enzyme] is the concentration of the target enzyme, and n_H_ is the Hill coefficient.

The limit of detection (LOD) was calculated based on the ratio of 3 times of standard deviation of the blank to the slope of linear regression at a low concentration range.^2^ All Statistical analyses were completed using GraphPad Prism 8.0.

The inhibition percentage for different inhibitors was calculated as follows:

$$Inhibition \left( \% \right)= 1-\frac{F_{inh}- F_{hOGG1 Transducer}}{F_{hOGG1 Transducer, Enzyme} - F_{hOGG1 Transducer}} \times100$$

Where *F_inh_* is the fluorescence signal of the hOGG1 DNA Transducer measured after the preincubation with different concentrations of inhibitors and hOGG1 (ng/ml) or Fpg (U/ml), *F_hOGG1 Transducer_*,_Enzyme_ represents the fluorescence signal of the hOGG1 DNA Transducer in the presence of hOGG1 (ng/ml) or Fpg (U/ml). In contrast, F_hOGG1 Transducer_ denotes the fluorescence signal in the absence of hOGG1 or Fpg. IC50 values were calculated using GraphPad Prism 8.0 software with a four-parameter logistic dose-response curve.

### DNA Sequences

All sequences were designed using NUPACK.^3^

**Table S1. UDG DNA Transducers used for UDG detection.**

| **Name** | **Sequence** | Figure |
| --- | --- | --- |
| UDG Transducer #0U | *AGATTTAGCCATGTGTAGAC***CAAA***GTCTACACATGGCTAAATCT* | 2B-2D |
| UDG Transducer #1U | *AGATTTAGCCATGTGTAGAC***CAAA***GTC(2-Deoxyuridine)ACACATGGCTAAATCT* | 2B-2D |
| UDG Transducer #2U | *AGATTTAGCCATGTGTAGAC***CAAA***GTCTACACA(2-Deoxyuridine)GGC(2-Deoxyuridine)AAATCT* | 2B-2D |
| UDG Transducer #3U | *AGATTTAGCCATGTGTAGAC***CAAA***GTCTACACA(2-Deoxyuridine)GGC(2-Deoxyuridine)AAA(2-Deoxyuridine)CT* | 2B-2D |
| UDG Transducer #4U | *AGATTTAGCCATGTGTAGAC***CAAA***GTC(2-Deoxyuridine)ACACA(2-Deoxyuridine)GGC(2-Deoxyuridine)AAA(2-Deoxyuridine)CT* | 2, 3, Supplementary Figures S3-S5 |
| UDG Transducer #14 | *AGCCATGTGTAGAC***CAAA***GTCTACACA(2-Deoxyuridine)GGCT* | 2E |
| UDG Transducer #16 | *TTAGCCATGTGTAGAC***CAAA***GTCTACACAUGGCTAA* | 2E |
| UDG Transducer #18 | *ATTTAGCCATGTGTAGAC***CAAA***GTCTACACA(2-Deoxyuridine)GGCTAAAT* | 2E |
| UDG Transducer #20 | *AGATTTAGCCATGTGTAGAC***CAAA***GTCTACACA(2-Deoxyuridine)GGCTAAATCT* | 2E |
| UDG Transducer #4U-FQ | (6-FAM)-*AGATTTAGCCATGTGTAGAC***CAAA***GTC(2-Deoxyuridine)ACACA(2-Deoxyuridine)GGC(2-Deoxyuridine)AAA(2-Deoxyuridine)CT*-(BHQ1) | 2F, Supplementary Figures S1, S2 |
| UDG Transducer #4AP-FQ | (6-FAM)-AGATTTAGCCATGTGTAGAC**CAAA***GTC()ACACA()GGC()AAA()CT*-(BHQ1) | Supplementary Figures S1, S2 |

Here, the *italic* sequences represent the duplex stem of DNA Transducers where the *underlined* bases are also complementary to the crRNA. The **bold** portion represents the loop of the DNA Transducer.

**Table S2. DNA Transducers used for hOGG1 detection.**

| **Name** | **Sequence** | Figure |
| --- | --- | --- |
| hOGG1 Transducer | *AGATTTAGCCATGTGTAGAC***CAAA***GTCTACACAT(8-oxoG)GCTAAATCT* | 4, 5, Supplementary Figures S6-S9 |
| hOGG1 Transducer w/o Goxo | *AGATTTAGCCATGTGTAGAC***CAAA***GTCTACACATGGCTAAATCT* | Supplementary Figures S6 |
| hOGG1 Transducer-FQ | *AGATTTAGCCATGTGTAGAC***CAAA***GTCTACACAT(8-oxoG)GCTAAATCT* | Supplementary Figure S7 |

In the above sequences, the **bold** portion represents the loop of the DNA Transducer. The *italic* sequences represent the duplex stem of DNA Transducers where the *underlined* bases complement the crRNA.

**Table S3. crRNA, FRET-based DNA reporter, ssDNA target**

| **Name** | **Sequence** | Figure |
| --- | --- | --- |
| crRNA | UAAUUUCUACUAAGUGUAGAU*GUCUACACAUGGCUAAAUCU* | 2-5, Supplementary Figures S3-S11 |
| FRET-based DNA reporter | (6-FAM)-CTCTCA**TTTTTTTTTT**AGAGAG-(BHQ 1) | 2-5, Supplementary Figures S3-S5, S7-S11 |
| ssDNA target | AGATTTAGCCATGTGTAGAC | Supplementary Figures S10 and S11 |

Here the *italic* bases represent the crRNA-targeting sequence. The **bold** and underlined sequences represent the loop and stem-forming portions of the FRET-based DNA reporter, respectively.

### Supplementary Figures

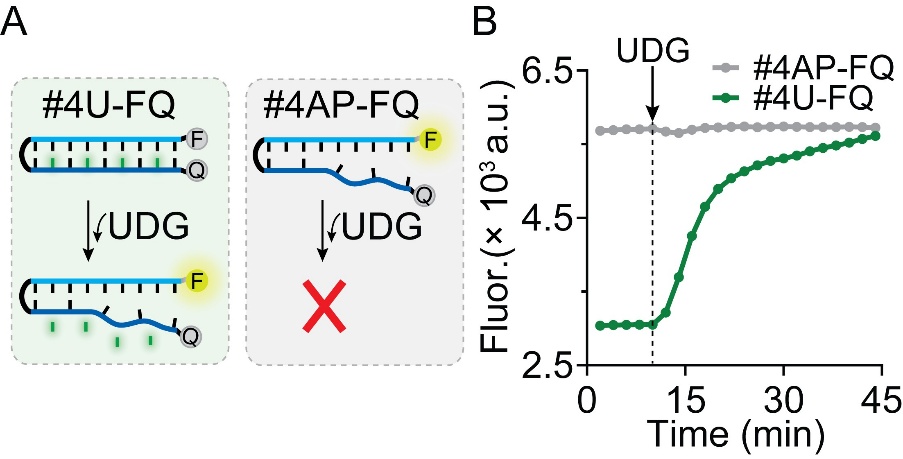

**Figure S1.** UDG repair activity on FRET-labelled DNA Transducers containing uracil lesions (UDG Transducer #4U-FQ) or AP sites (UDG Transducer #4AP-FQ). (A) Scheme of the reaction showing that only the DNA Transducer containing U bases can be detected by UDG. (B) Time-course fluorescence experiments of UDG activity using the two FRET-labelled DNA Transducers showing that an increase in the FAM emission due to the destabilization of the hairpin is observed only for the DNA Transducer #4U-FQ. As expected the transducer containing four abasic sites is already unfolded and does not interact with UDG. Experiments were performed at 37 °C in a buffer solution (10 mM Tris-HCl, 50 mM NaCl, 10 mM MgCl₂, and 0.1 mg/ml BSA, at pH 7.9) containing 500 nM concentration of FRET-labelled Transducer by adding 10 U/ml of UDG.

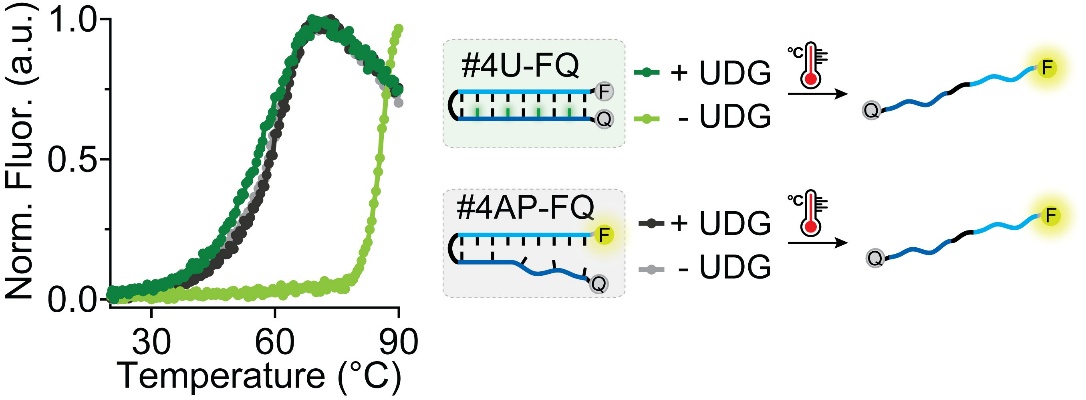

**Figure S2.** Melting curves of FRET-labeled UDG Transducers containing either uracil lesions (UDG Transducer #4U-FQ) or abasic (AP) sites (UDG Transducer #4AP-FQ) were recorded in the presence and absence of UDG. The melting profile of the transducer with four AP sites remained unchanged upon UDG treatment, indicating no further processing. In contrast, the UDG Transducer #4U-FQ exhibited a reduced melting temperature (Tm = 59.6 °C) following UDG activity, aligning with the melting profile of the #4AP-FQ construct. These results confirm the enzymatic conversion of uracil lesions into AP sites by UDG. Fluorescence melting curves were obtained by heating a buffer solution containing 500 nM concentration of FRET-labelled Transducer, either in the presence or absence of UDG, from 20 °C to 90 °C at a rate of 1 °C min^–1^ and monitoring fluorescence at every 0.5 °C, setting λ_ex._= 490 nm and λ_em_. = 520 nm.

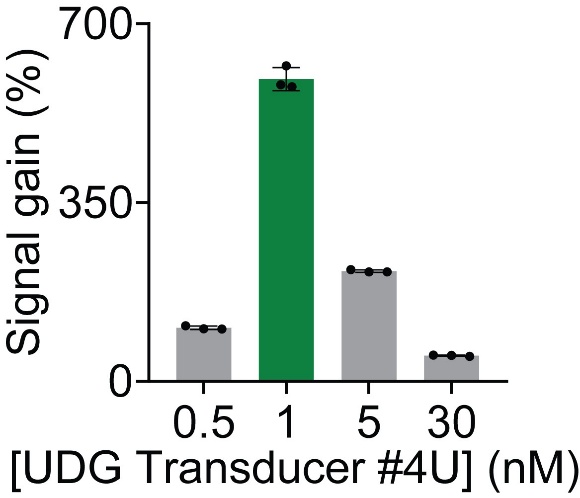

**Figure S3.** Signal gain (%) calculated after 15 min of UDG activity and different concentrations of UDG DNA Transducer #4U. Experiments were conducted at 37 °C in a buffer solution (10 mM Tris-HCl, 50 mM NaCl, 10 mM MgCl₂, and 0.1 mg/ml BSA, at pH 7.9) containing different concentrations of DNA Transducer and 0.5 U/ml of UDG by adding the Cas12a reaction mix (500 nM of FRET-based DNA reporter and 20 nM of Cas12a/crRNA complex). Error bars represent the deviation from three independent experiments.

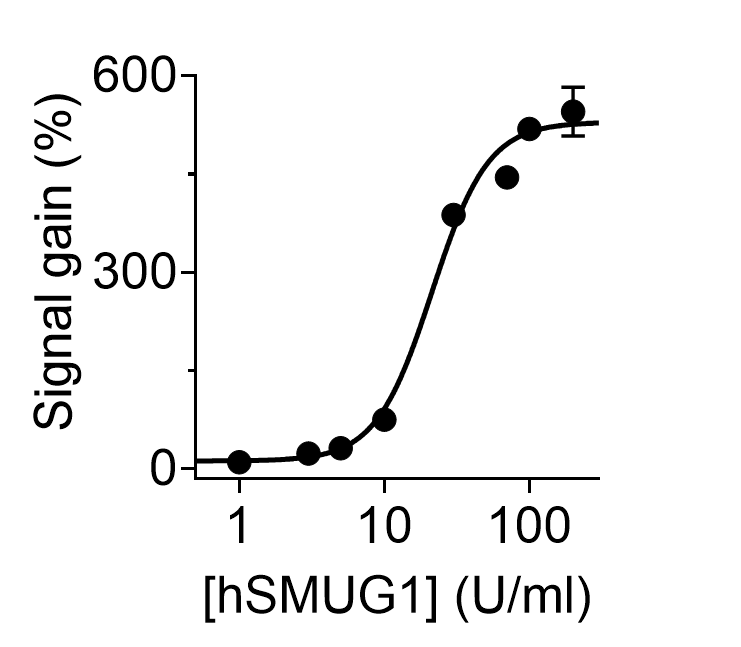

**Figure S4.** Plots of signal gain (%) as a function of hSMUG1 concentration obtained after 15 minutes of cleavage reaction. Experiments were performed at 37 °C in a buffer solution (10 mM Tris-HCl, 50 mM NaCl, 10 mM MgCl₂, and 0.1 mg/ml BSA, at pH 7.9) containing 1 nM concentration of DNA Transducer #4U and Cas12a reaction mix (500 nM of FRET-based DNA reporter and 20 nM of Cas12a/crRNA complex) by adding an increasing concentration of hSMUG1. Error bars represent the deviation from three independent experiments.

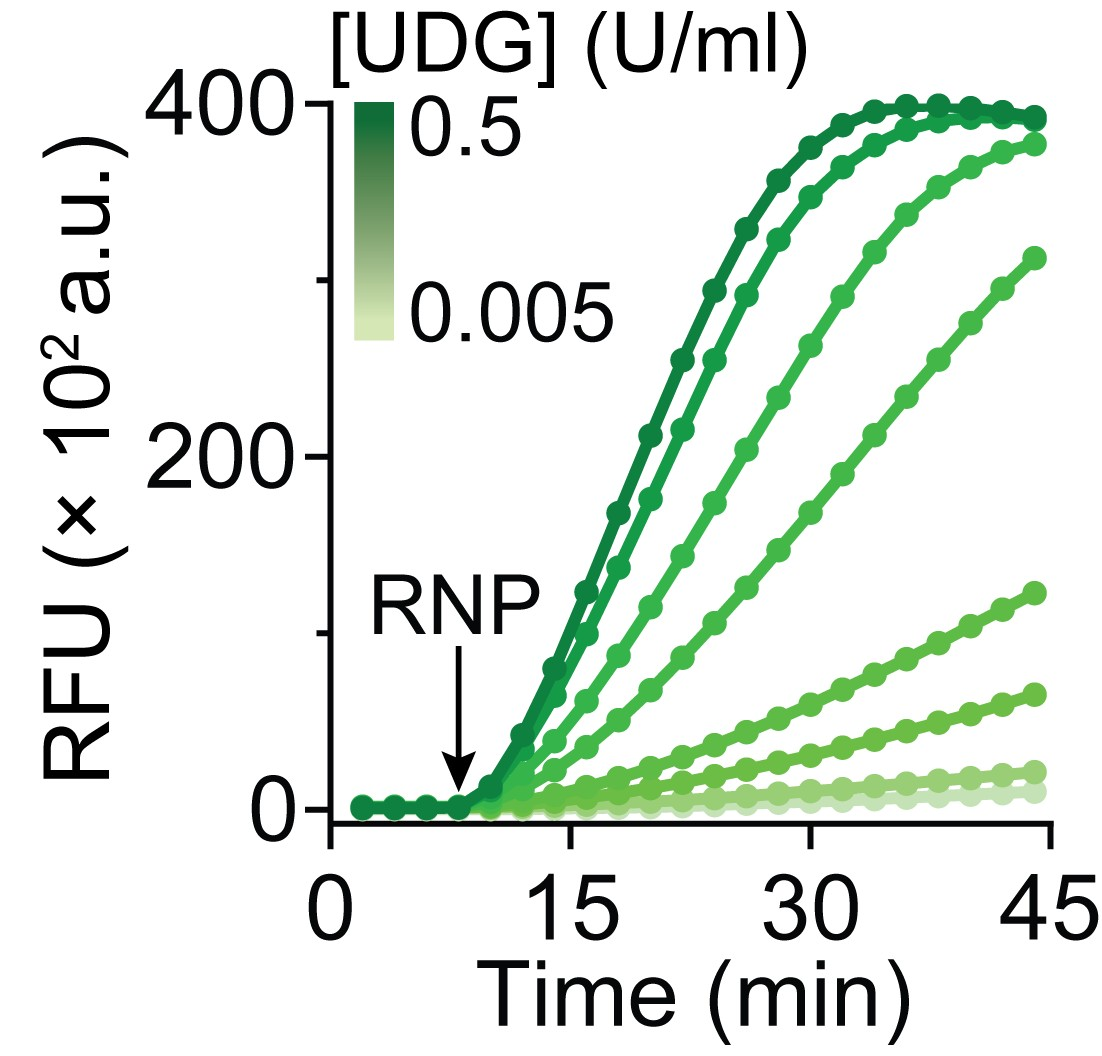

**Figure S5.** Kinetic traces obtained at increasing concentrations of spiked UDG into inactivated HEK293 T cell lysate. Experiments were performed at 37 °C in 10 µg/ml lysate sample containing 1 nM #4U, and Cas12a reaction mix (500 nM of FRET-based DNA reporter and 20 nM of Cas12a/crRNA) by adding increasing concentrations of UDG.

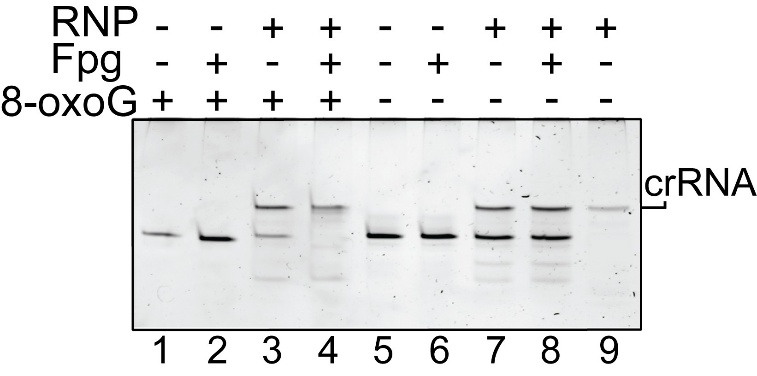

**Figure S6.** Denaturing PAGE analysis of Cas12a *cis*-cleavage activity in the presence and absence of Fpg, using hOGG1 Transducers that either contain (hOGG1 Transducer, lanes 1 to 4) or lack (hOGG1 Transducer w/o Goxo, lanes 5 to 8) the damage base 8-oxoG.

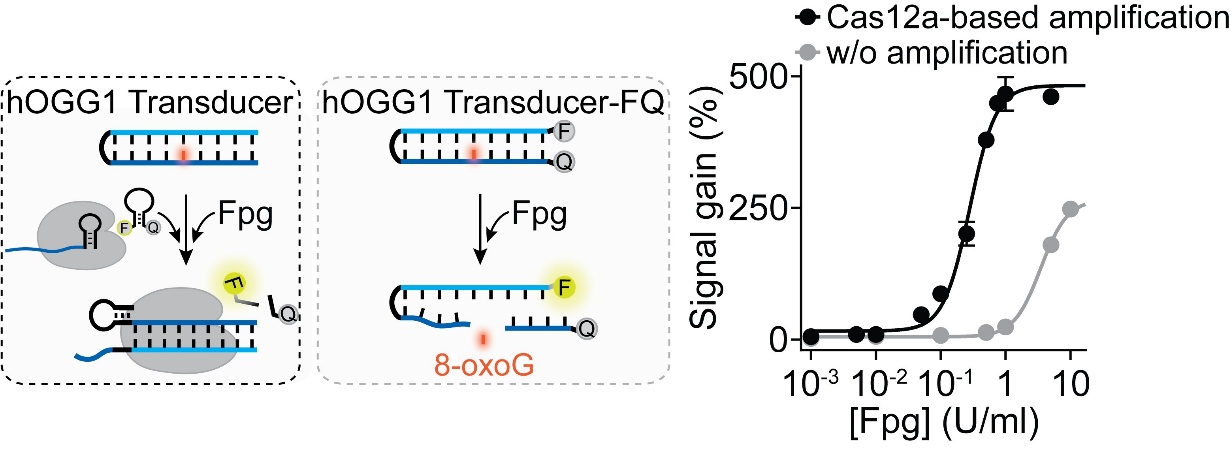

**Figure S7**. Comparison between the dose-response curves obtained using the Cas12a-based Fpg activity platform and the direct FRET-based assay using FAM-BHQ1 labelled hOGG1 Transducer (hOGG1 Transducer -FQ, 500 nM) after 15 min of reaction with the enzyme. Our Cas12a UDG activity assay was performed using hOGG1 Transducer (1 nM) and the Cas12a reaction mix (500 nM of FRET-based DNA reporter and 20 nM of Cas12a/crRNA complex) in the presence of increasing concentrations of Fpg. Error bars represent the deviation from three independent experiments. Both the assays were performed in a buffer solution (10 mM Tris-HCl, 50 mM NaCl, 10 mM MgCl₂, and 0.1 mg/ml BSA, at pH 7.9) at 37°C.

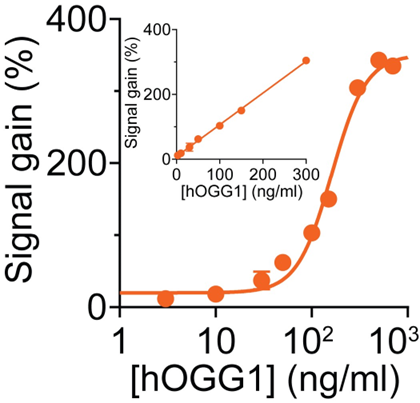

**Figure S8**. Calibration curve of hOGG1 activity performed in 10 µg/ml lysate sample. The inset shows the linear correlation between signal gain (%) values and hOGG1 concentrations in the range of 3 to 300 ng/ml. Experiments were performed at 37 °C in a 10 µg/ml lysate sample containing hOGG1 Transducer (1 nM) and the Cas12a reaction mix (500 nM FRET-based DNA reporter and 20 nM Cas12a/crRNA complex), by adding increasing concentrations of hOGG1 and collecting the signal after 15 minutes of reaction. Error bars represent the deviation from three independent experiments.

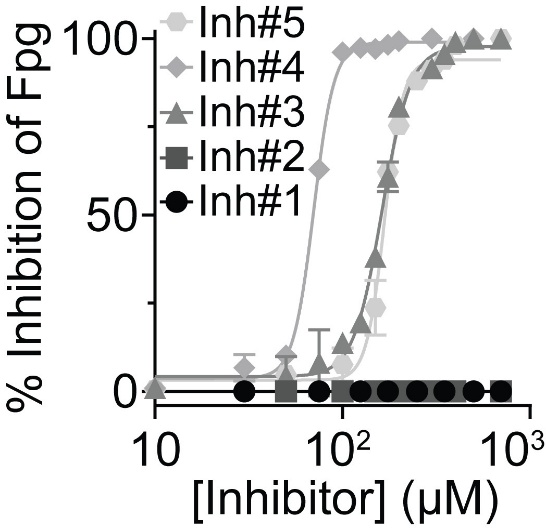

**Figure S9**. Plot of % inhibition of Fpg activity at various concentrations of different inhibitors. Experiments were performed by adding a Cas12a reaction mix (500 nM FRET-based DNA reporter, 20 nM Cas12a/crRNA complex, and 1 nM of hOGG1 Transducer) to a buffer solution (10 mM Tris-HCl, 50 mM NaCl, 10 mM MgCl₂, and 0.1 mg/ml BSA, at pH 7.9) containing Fpg (0.75 U/ml), preincubated with various concentrations of inhibitors at 37 °C. Inh#1 to #5 represent O8, SU0268, 2-thioxanthine, Mercapto-6,8-purinediol, and 8-mercaptoadenine, respectively.

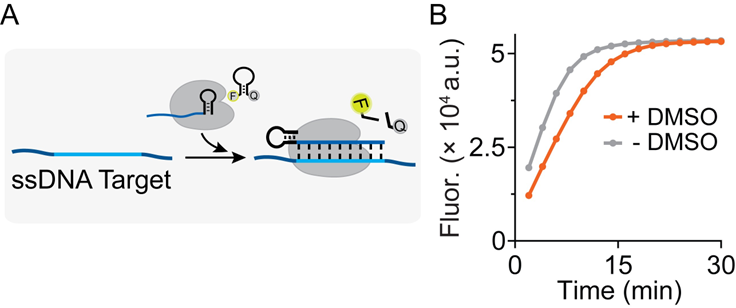

**Figure S10**. Effect of 10% DMSO on *trans*-cleavage activity of Cas12a/crRNA complex. A) Scheme of reaction. B) Kinetic traces of Cas12a *trans*-cleavage activity on a complementary ssDNA target in the absence (gray trace) or presence (orange trace) of 10% DMSO. Experiments were carried out at 37 °C by adding a Cas12a/crRNA complex (20 nM) to a buffer solution containing 1 nM ssDNA target and 500 nM FRET-based DNA reporter, with or without 10% DMSO.

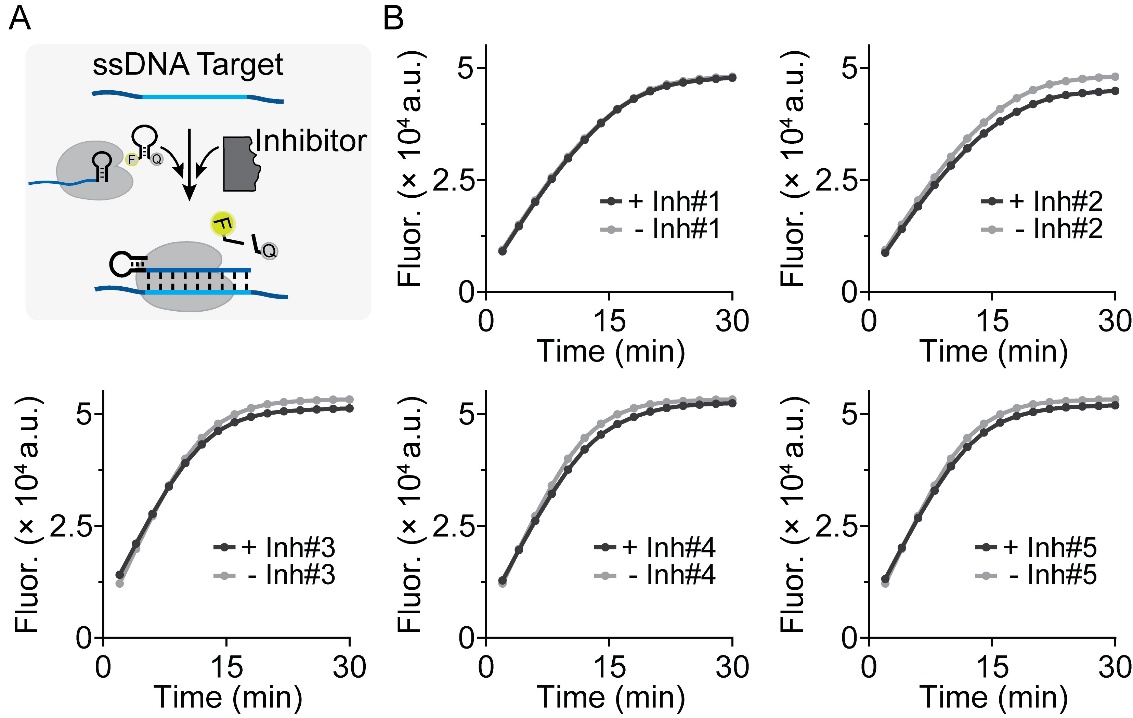

**Figure S11**. Kinetic traces of *trans*-cleavage activity using a ssDNA input in the absence (gray trace) or presence (orange trace) of different inhibitors. Data reported confirm that all the tested inhibitors do not affect the *trans*-cleavage activity of Cas12a. Experiments were carried out at 37 °C by adding a Cas12a/crRNA complex (20 nM) to a buffer solution containing 1 nM ssDNA target, and 500 nM FRET-based DNA reporter with or without 350 µM inhibitors. Inh#1 to #5 represent O8, SU0268, 2-thioxanthine, Mercapto-6,8-purinediol, and 8-mercaptoadenine, respectively.
